## SUPPLEMENTARY DATA for "Three ParA dimers cooperatively assemble on type Ia partition promoters"

**This PDF file includes:**

### **Supplementary Figure S1. ParA_F_ purification steps and specific DNA binding activity.**

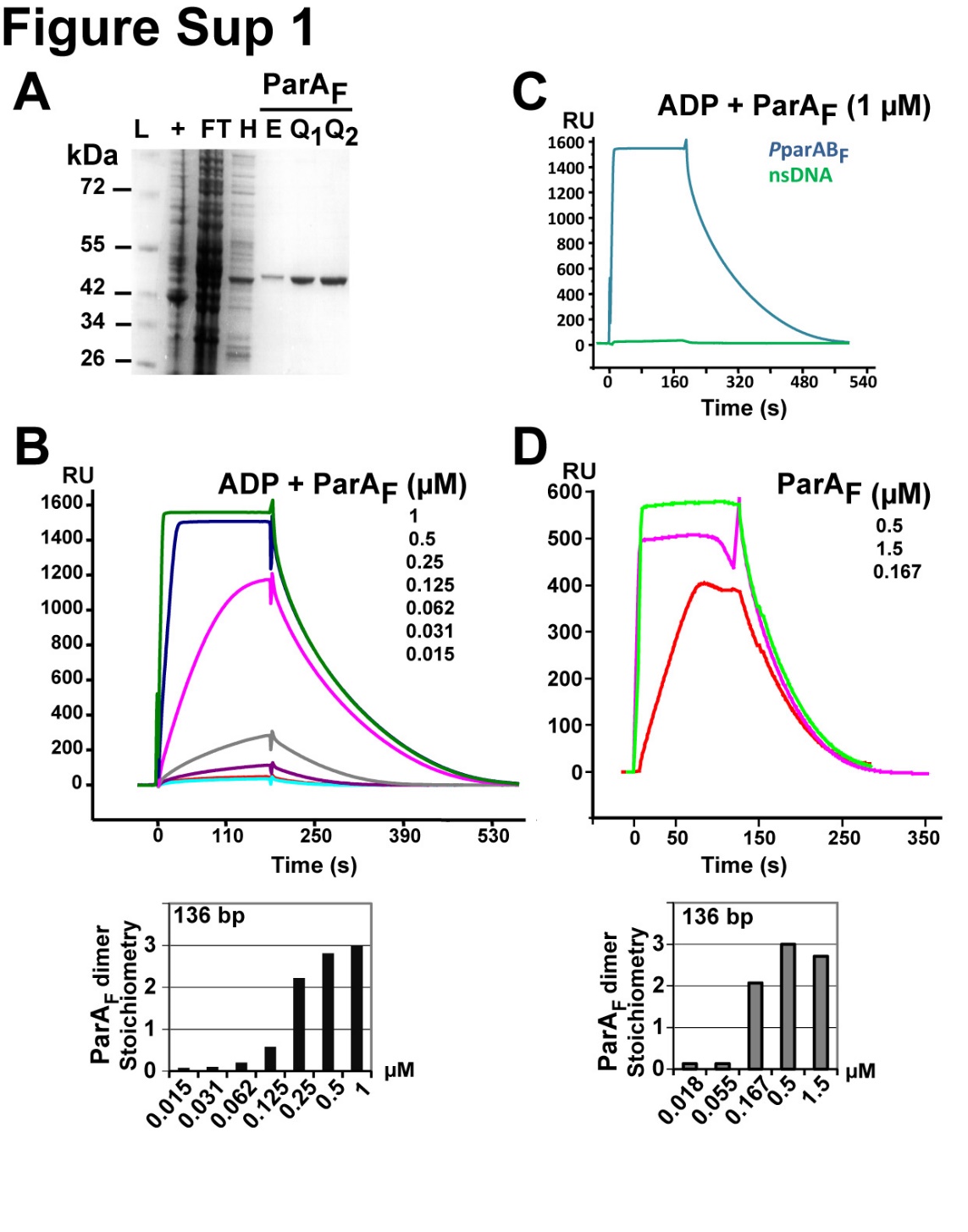

**A.** Purification steps of ParA_F_. The protein fractions during the purification procedure were analyzed on a 4-20% SDS-PAGE and colored with Instant Blue (Euromedex). Samples were crude extract from induced culture (+), flow through of the heparin column (FT), heparin fraction (H), exclusion fraction of Superdex 200 (E), monoQ fractions (Q1-Q2). Molecular weight ladder (L) is indicated on the left in kDa.

**B.** SPR analysis of ParA_F_ binding to a 136-bp *P*parAB_F_ DNA fragments in presence of 1 mM ADP. Upper panel: 530 RU of biotinylated DNA was immobilized on streptavidin SA chip (Biacore 3000). Increasing concentration of ParA_F_ (from 0.015 to 1 µM) was injected (time 0) at 10 µL/min for 180 s before dissociation. Lower panel: stoichiometry of ParA_F_ dimers bound per *P*parAB_F_ DNA fragment, calculated as in Figure 2B. Three ParA_F_ dimers are bound per DNA fragment at saturation.

**C.** ParA_F_ binds specifically to the promoter region in the presence of ADP. SPR analysis (Biacore 3000) of ParA_F_ binding to a 136-bp DNA fragments containing *P*parAB_F_ (blue) or nsDNA (green) performed as in Figure 2B except for the presence of 1 mM ADP in the reaction.

**D.** SPR analysis as in (B) using a Biacore X100 with 200 RU of 136-bp *P*parAB_F_ fragment immobilized. ParA_F_ proteins (from 0.018 to 1.5µM) were injected at 30 µL/min. The three higher concentrations, 0.17, 0.5 and 1.5 µM are displayed in red, green and purple, respectively. Lower panel: stoichiometry of ParA_F_ dimers bound per *P*parAB_F_ DNA fragment, calculated as in Figure 2B. Three ParA_F_ dimers are bound per DNA at saturation without ADP.

### **Supplementary Figure S2. Sequences of the oligonucleotides used in this study.**

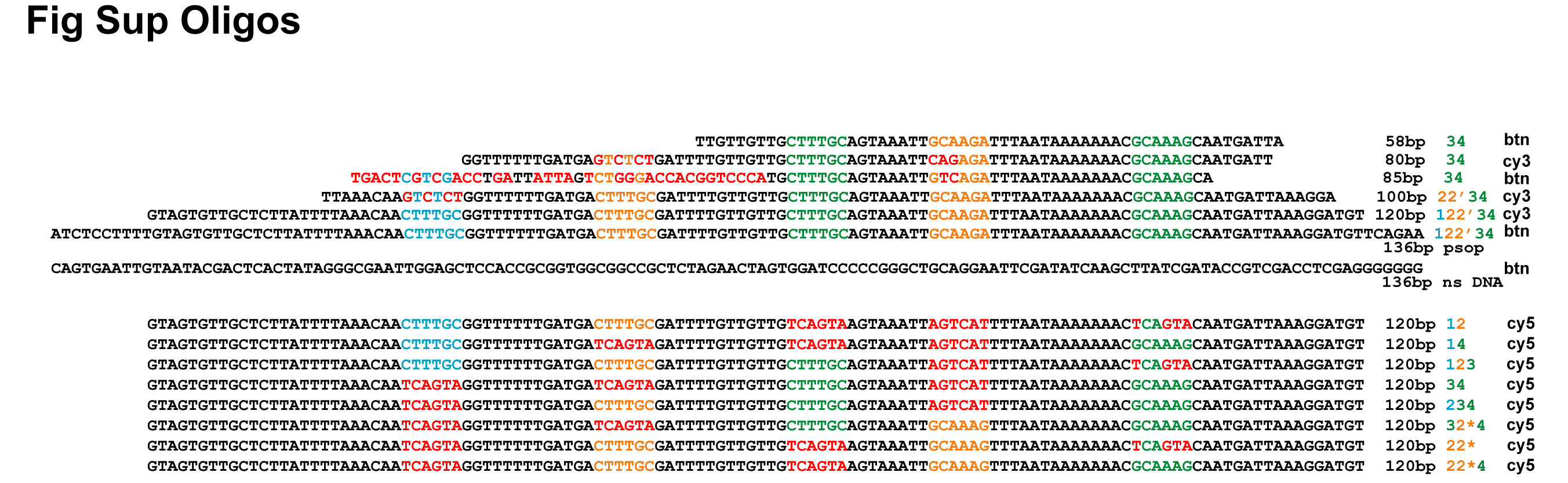

The indicated oligonucleotides are annealed with their complementary oligonucleotides to generate duplex DNA probes. Note that each DNA fragment is designated by its length in base pair (bp). The presence of the binding motifs is indicated by their numbering at the end of the line. Cy3 or biotin (btn) modifications at the 5’ extremity of the indicated oligonucleotides are also indicated. Motifs are color coded as in Figure 5C.

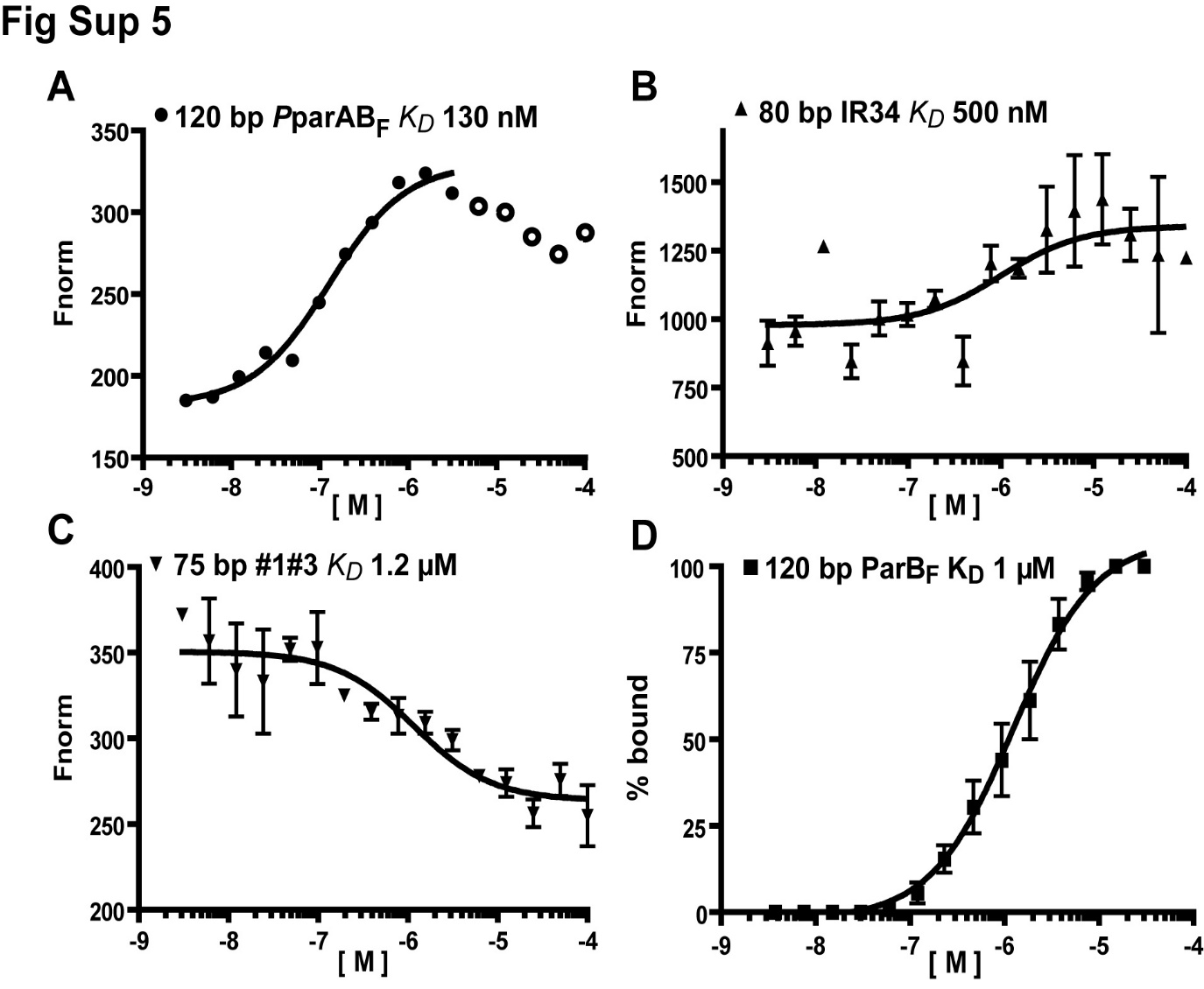

### **Supplementary Figure S3. Weak DNA binding of ParA_F_ on DNA fragment smaller than 80 bp and non-specific DNA binding of ParB_F_.**

Quantification of the DNA binding affinities of ParA_F_ (A-C) and ParB_F_ (D) to various DNA substrates measured by fluorescence (A-C) or EMSA (D). The resulting curves were fitted using non-linear regression by Prism to calculate an apparent *K_D_*. Experiments have been reproduced three times except for (A).

**A.** A 120-bp Cy3 labelled DNA probe (25 nM) containing *P*parAB_F_ was titrated with 3.6 nM to 100 µM ParA_F_. The fluorescence measurements lead to an apparent *K_D_* of 130 nM. Decreasing signals above 2µM was due to aggregation, detected by denaturation measurement within the assays.

**B.** A 80-bp Cy3 labelled DNA probe (15 nM) containing IR3-4 was titrated with 3.6 nM to 100 µM ParA_F_. The fluorescence measurements lead to an apparent *K_D_* of 500 ± 325 nM.

**C.** A 75-bp Cy3 labelled DNA probe (15 nM) containing the binding motifs #1 and #3 was titrated with 3.6 nM to 100 µM ParA_F_. The fluorescence measurements lead to an apparent *K_D_* of 1200 ± 700 nM.

**D.** A 120-bp Cy3 labelled DNA probe (15 nM) was titrated with 3.7 nM to 30 µM ParB_F_ and analyzed in EMSA. The quantification of the free versus retarded DNA leads to an apparent *K_D_* of 1 ± 0.2 µM, in agreement with previous measurements (Ah-Seng et al., 2009).

### **Supplementary Figure S4. Secondary structure analysis of ParA_F_.**

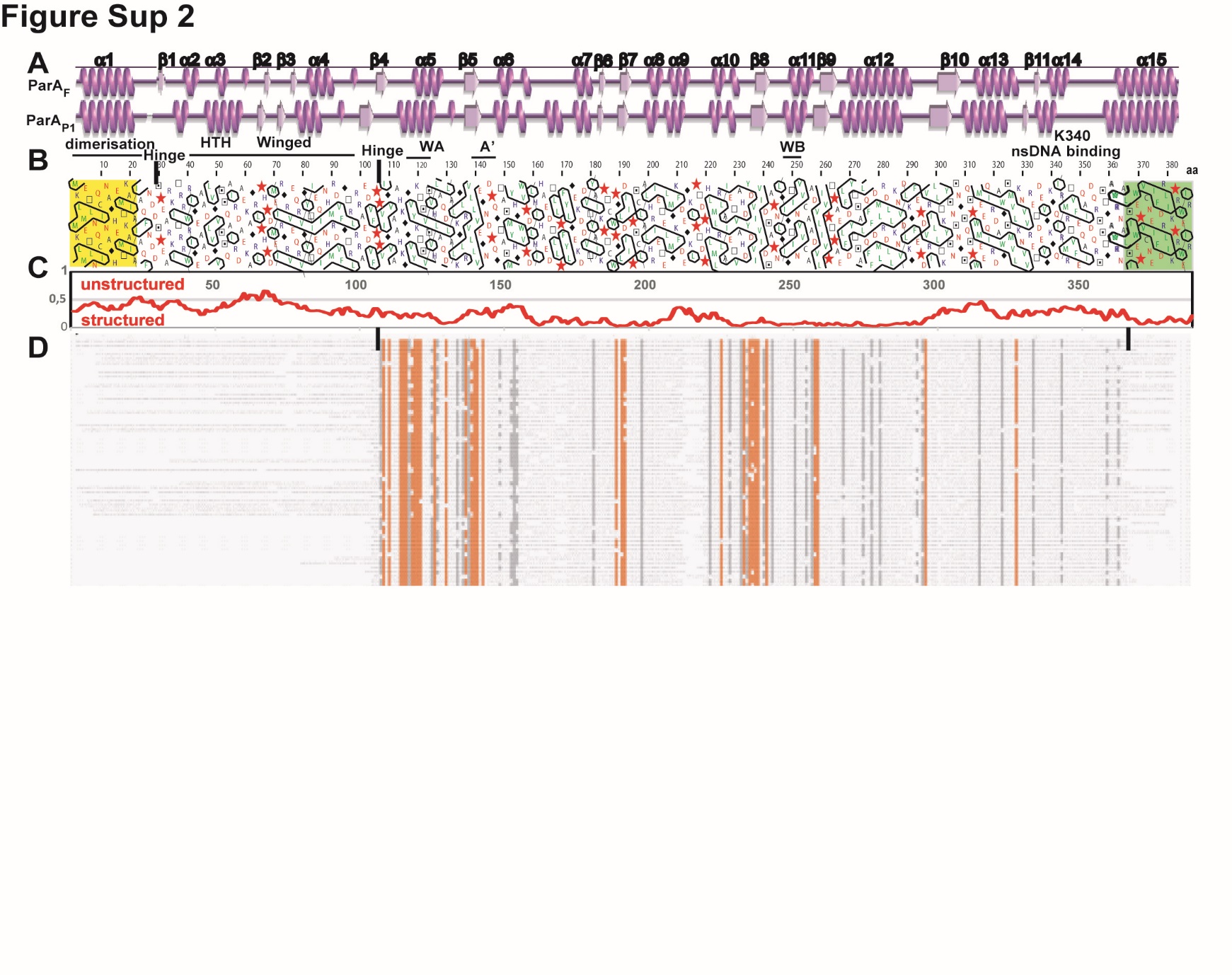
**A.** Comparison of secondary structure assignment of 3D model of ParA_F_ and X-ray structure of ADP ParA_P1_ (3ez2) generated by PDBsum (de Beer et al., 2014). The α-helices and β-strands are numbered according to ParA_F_.

**B**. Hydrophobic cluster analysis plot of ParA_F_ protein (Callebaut et al., 1997). The numbering of residues is labelled on top. Key amino acids are highlighted by a symbol, proline (star), glycine (diamond), threonine (empty square) and serine (dotted square). The different domains and motifs are labelled as in Figure 4A. First and last α-helices are colored yellow and green respectively. **C**. Analysis of intrinsically unstructured/disordered domains of ParA_F_ protein performed using IUPred (Dosztányi et al., 2005). **D.** Domains conservation of ParA_F_ proteins. A Psi blast alignment profile was automatically constructed from a BLAST search using ParA_F_ (UniProtP 62556) as query (Altschul et al., 1997). Sixty sequences representing the archetypes of types I are represented over a thousand sequences that were recovered and aligned. Both subtypes Ia and Ib share the conserved central core corresponding to the Walker boxe (Gerdes et al., 2000). The conserved residues are colored in orange. Black verticals bars represent the limit between the non-conserved and the most conserved regions. The latter corresponds to the Walker-box domain.

### **Supplementary Figure S5. *P*parAB_F_ homologous sequence.**

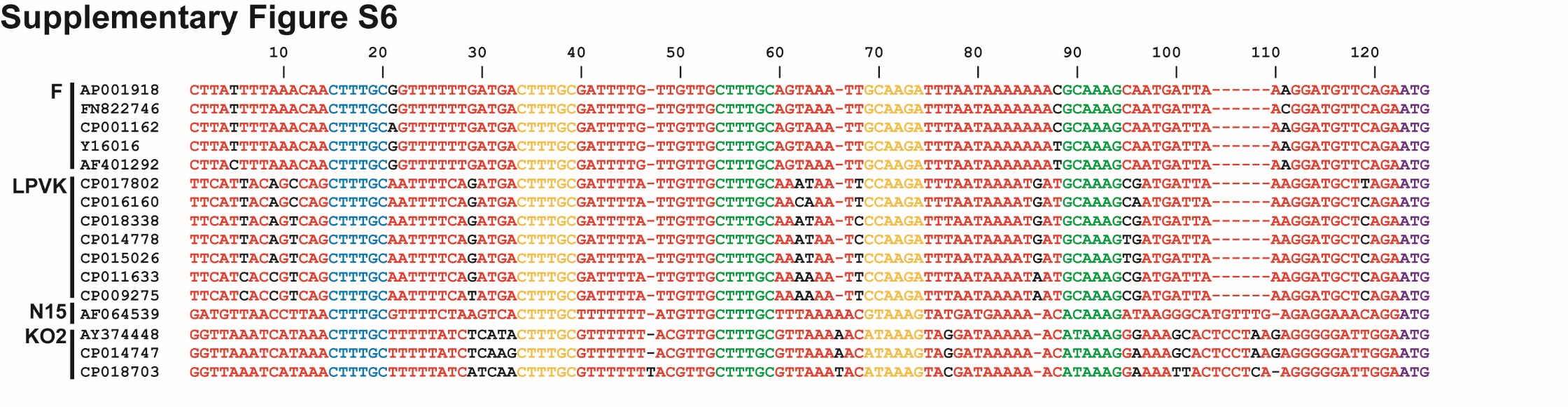

Homologous sequences of *P*parAB_F,_ *P*parAB_LPVK,_ *P*parAB_N15,_ and *P*parAB_KO2_ were searched by Blastn analyses. Only the sequences presenting at least one nucleotide difference are displayed. The first sequence of each group (indicated on the left) corresponds to the promoter used for the Blastn search, with the nucleotide variation within a group labelled in black. Sequences closely related to *P*parAB_F_ are: AP001918 (*E. coli* K-12 plasmid F), FN822746 *(E. coli* ETEC 1392/75 plasmid p557), CP001162 (*E. coli* Vir68 plasmid pVir68), Y16016 (*E. coli* plasmid pO157), AF401292 *(E. coli* O157:H- plasmid pSFO157).

Sequences closely related to *P*parAB_LPVK_ are: CP017802 (*Raoultella ornithinolytica*), CP016160 (*Klebsiella pneumoniae*), CP018338 (*K. pneumoniae* isolate Kp_Goe_154414 plasmid pKp_Goe_414-2), CP014778 (*Pluralibacter gergoviae* strain FB2 plasmid pFB2.3), CP015026 (*K. pneumoniae* strain Kpn223 plasmid pKPN-065), CP011633 (*Klebsiella oxytoca* strain CAV1374 plasmid pCAV1374-150), CP009275 (*K.  variicola* strain DX120E plasmid pKV1). No closely related sequence to *P*parAB_N15_ (AF064539) was found. Sequences closely related to *P*parAB_KO2_ are: AY374448 (*Bacteriophage phiKO2*), CP014747 (*Enterobacter aerogenes* strain FDAARGOS_139 plasmid), CP018703 (*K. pneumoniae* strain Kp_Goe_827024 plasmid pKp_Goe_024-5). ParA_F_ binding motifs are colored as in Figure 5C. The ATG start codons at the end of the sequence are colored in purple.

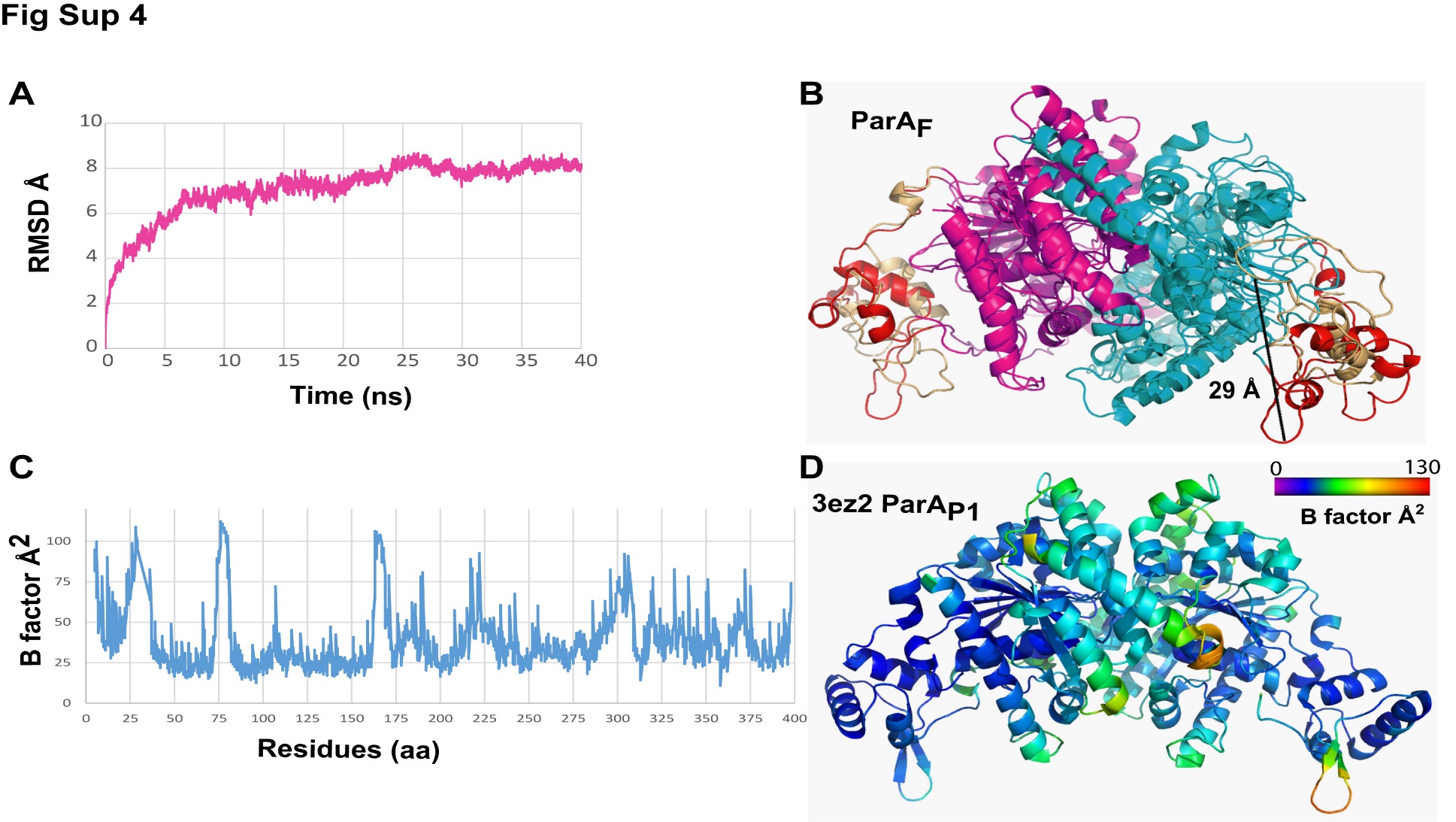

### **Supplementary Figure S6. ParA_F_ and ParA_P1_ dimers undergo large conformation changes in the wing domain.**

**A**. Quantitative measurement of ParA_F_ movements during MD simulations. Root mean squared deviation of ParA_F_ backbone atoms of ParA_F_ monomer is displayed as a function of time during 40 ns simulation of in explicit water. **B**. Backbone superposition of the starting model of ParA_F_ dimer at 0 ns and the snapshot after 40 ns MD simulation (monomers are colored pink and cyan). Winged-HTH region of two states (0 and 40 ns) are colored in red and salmon, respectively. Distance between the arginine 75 C-alpha atoms in initial and 40 ns structure is shown in black. **C**. Plot of the ParA_P1_ residues B-factors reported in the 3ez2 PDB files (Dunham et al., 2009). **D.** Schematics of ParA_P1_ X-ray structure. The fold is shown in cartoon and colored-coded according to B-factors plotted in (C).

### **Supplementary Table S1. Summary of the binding and kinetic parameters determined by SPR analyses.**

| ***K_D_*** | ***ka* M^-1^ s^-1^** | ***kd* s^-1^** | **Chi^2^** | **R_max_** | **^RU immobilized^** | **^Flow µL/min^** | **^Apparatus^** |
| --- | --- | --- | --- | --- | --- | --- | --- |
| 472 nM*^a^* | 1.9 10^4^ | 0,0092 | 490 | 2300 | 530 | 10 | Biacore3000 |
| 640 nM*^b^* | 1,39 10^4^ | 0,0089 | 220 | 2000 | 530 | 10 | Biacore3000 |
| 450 nM*^c^* | 6,89 10^4^ | 0,0318 | 880 | 850 | 300 | 20 | Biacore3000 |
| 70 nM*^d^* | 3,87 10^5^ | 0,0274 | 1000 | 650 | 200 | 30 | BiacoreX100 |

The binding and kinetics parameters from the SPR analyses assaying the ParA_F_-*P*parABS_F_ interaction described in the main text and Supplementary materials are summarized along with the experimental conditions and apparatus used for each analysis. All experiments were performed on a 136-bp DNA fragments containing *P*parAB_F_ region. Note that the Chi^2^ values for the experiments performed with conditions ^c^ and ^d^ are close to the Rmax values indicating that the accuracy of these experiments is low. As mentioned in the main text, these variations depend mainly on the tendency of ParA_F_ to self-aggregate and therefore the kinetics parameters have to be taken as an indication of the interaction.

*^a^* Data sets in which ParA_F_ concentrations vary from 0.015 to 1 µM.

*^b^* Data sets performed in the presence of 1 mM ADP from Figure S1B.

*^c^* Data sets from Figure 2A.

*^d^* Data sets from Supplementary Figure S1D.

### **Suplementary references**

Ah-Seng, Y., Lopez, F., Pasta, F., Lane, D., and Bouet, J.-Y. (2009). Dual Role of DNA in Regulating ATP Hydrolysis by the SopA Partition Protein. J. Biol. Chem. *284*, 30067–30075.

Altschul, S.F., Madden, T.L., Schäffer, A.A., Zhang, J., Zhang, Z., Miller, W., and Lipman, D.J. (1997). Gapped BLAST and PSI-BLAST: a new generation of protein database search programs. Nucleic Acids Res. *25*, 3389.

de Beer, T.A.P., Berka, K., Thornton, J.M., and Laskowski, R.A. (2014). PDBsum additions. Nucleic Acids Res. *42*, D292–D296.

Callebaut, I., Courvalin, J.C., Worman, H.J., and Mornon, J.P. (1997). Hydrophobic cluster analysis reveals a third chromodomain in the Tetrahymena Pdd1p protein of the chromo superfamily. Biochem Biophys Res Commun *235*, 103–107.

Dosztányi, Z., Csizmok, V., Tompa, P., and Simon, I. (2005). IUPred: web server for the prediction of intrinsically unstructured regions of proteins based on estimated energy content. Bioinformatics *21*, 3433–3434.
